# Interpretation of exercise-induced changes in human skeletal muscle mRNA expression depends on the timing of the post-exercise biopsies

**DOI:** 10.1101/2020.08.05.239038

**Authors:** Jujiao Kuang, Cian McGinley, Matthew J-C Lee, Nicholas J Saner, Andrew Garnham, David J Bishop

## Abstract

**Aim:** Exercise elicits a range of adaptive responses in skeletal muscle that include changes in mRNA expression. To better understand the health benefits of exercise training, it is essential to investigate the underlying molecular mechanisms of skeletal muscle adaptations to exercise. However, most studies have assessed the molecular events at a few convenient time points within a short time frame post exercise, and the variations of gene expression kinetics have not been addressed systematically.

**Method:** Muscle biopsies were collected from nine participants at baseline and six time points (0, 3, 9, 24, 48, and 72 h) following a session of high-intensity interval exercise. We assessed the mRNA content of 23 gene isoforms from the muscle samples.

**Result:** The temporal patterns of target gene expression were highly variable and the mRNA contents detected were largely dependent on the muscle sample timing. The maximal levels of mRNA content of all tested target genes were observed between 3 to 48 h post exercise.

**Conclusion:** Our findings highlight a critical gap in knowledge regarding the molecular response to exercise, where the use of a few time points within a short period after exercise has led to an incomplete understanding of the molecular responses to exercise. The timing of muscle sampling for individual studies needs to be carefully chosen based on existing literature and preliminary analysis of the molecular targets of interest. We propose that a comprehensive time-course analysis on the exercise-induced transcriptional response in humans will significantly benefit the field of exercise molecular biology.

## INTRODUCTION

Exercise is a powerful stimulus affecting skeletal muscle, leading to improvements in cardiovascular function, mitochondrial content and function, and whole-body metabolism ^1-4^. The molecular bases of skeletal muscle adaptations to exercise fundamentally involve increased protein content and enzyme activity, mediated by an array of pre- and post-transcriptional processes, as well as translational and post-translational control ^5,6^. From the onset of exercise, muscle contraction can induce disruptions to muscle homeostasis, including mechanical stress, calcium release, ATP turnover, changes to mitochondrial redox state, and reactive oxygen species (ROS) production. These cellular perturbations lead to the activation of signaling molecules, which activate a range of transcription factors and coactivators, such as peroxisome proliferator-activated receptor gamma coactivator 1α (PGC-1α) and p53 ^6^. In turn, changes in these and other proteins help to coordinate the transcription of genes associated with mitochondrial biogenesis, fat metabolism, and glucose metabolism ^6^.

It has been proposed that training-induced adaptions are due to the cumulative effect of each single exercise session, and that investigating exercise-induced changes in mRNA after a single exercise session can provide important information about the likely adaptations to repeated exercise sessions (i.e., exercise training) ^5,7^. Using both quantitative real-time PCR (qPCR) and whole-genome analysis, several studies have provided a better understanding of the transcriptional response to exercise ^8-22^. However, few studies have examined more than two or three time points post-exercise. While some studies have employed more sampling time points, most of the time points were within the first 24 h after exercise ^23-25^, and only a few studies have examined the mRNA expression beyond the first day ^26,27^. Thus, while there is a general understanding that exercise-induced changes in mRNA expression are time-dependent, few studies have investigated changes in exercise-induced gene expression at multiple time points post exercise.

The absence of a strong justification for the choice of post-exercise biopsy times has important implications for our understanding of molecular adaptations to exercise. For example, while Scribbans *et al.* ^28^ reported that there was not a systematic upregulation of nuclear and mitochondrial genes 3 h post exercise, they also noted their chosen biopsy time point may have failed to capture a systemic upregulation that occurred later in the post-exercise period. Similarly, the lack of increase in *p53* mRNA after exercise has been interpreted as evidence that post-translational modification is more important in regulating protein levels of p53 ^29^, although it is possible that exercise-induced changes in *p53* mRNA have been missed by the biopsy time points chosen to date. Thus, it is clear that the choice of post-exercise biopsy times can influence the interpretation of the transcriptional response to exercise ^23,27^.

The purpose of this research was to investigate the on- and off-kinetics of commonly assessed, exercise-responsive genes after a single session of endurance-based exercise. We assessed the mRNA expression of key transcription factors associated with exercise-induced mitochondrial biogenesis (PGC-1α and p53), as well as other genes that have potential roles in mitochondrial and metabolic adaptions to exercise. We hypothesised that different genes would elicit different temporal patterns of expression. The results have helped to highlight the importance of appropriate muscle sample timing and to provide recommendations for designing future studies examining molecular responses to exercise.

## RESULTS

### Dynamic gene expression response to exercise in human skeletal muscle

The influence of biopsy timing on mRNA content was examined following a single session of HIIE. We measured the mRNA content of 22 genes (23 isoforms) that have been implicated in the adaptive response to exercise (Table 1). The time point that elicited the highest (or lowest) mRNA content of each gene varied from 3 h to 72 h. To illustrate the distinct time course of mRNA expression in response to exercise, as well as the research interest of our group (mitochondrial adaptive responses to exercise), we chose to focus on nine genes (ten isoforms) that are related to mitochondrial adaptations in response to exercise, and which showed different time points for peak expression (Table 2 and Figure 1).

**Table 1.**
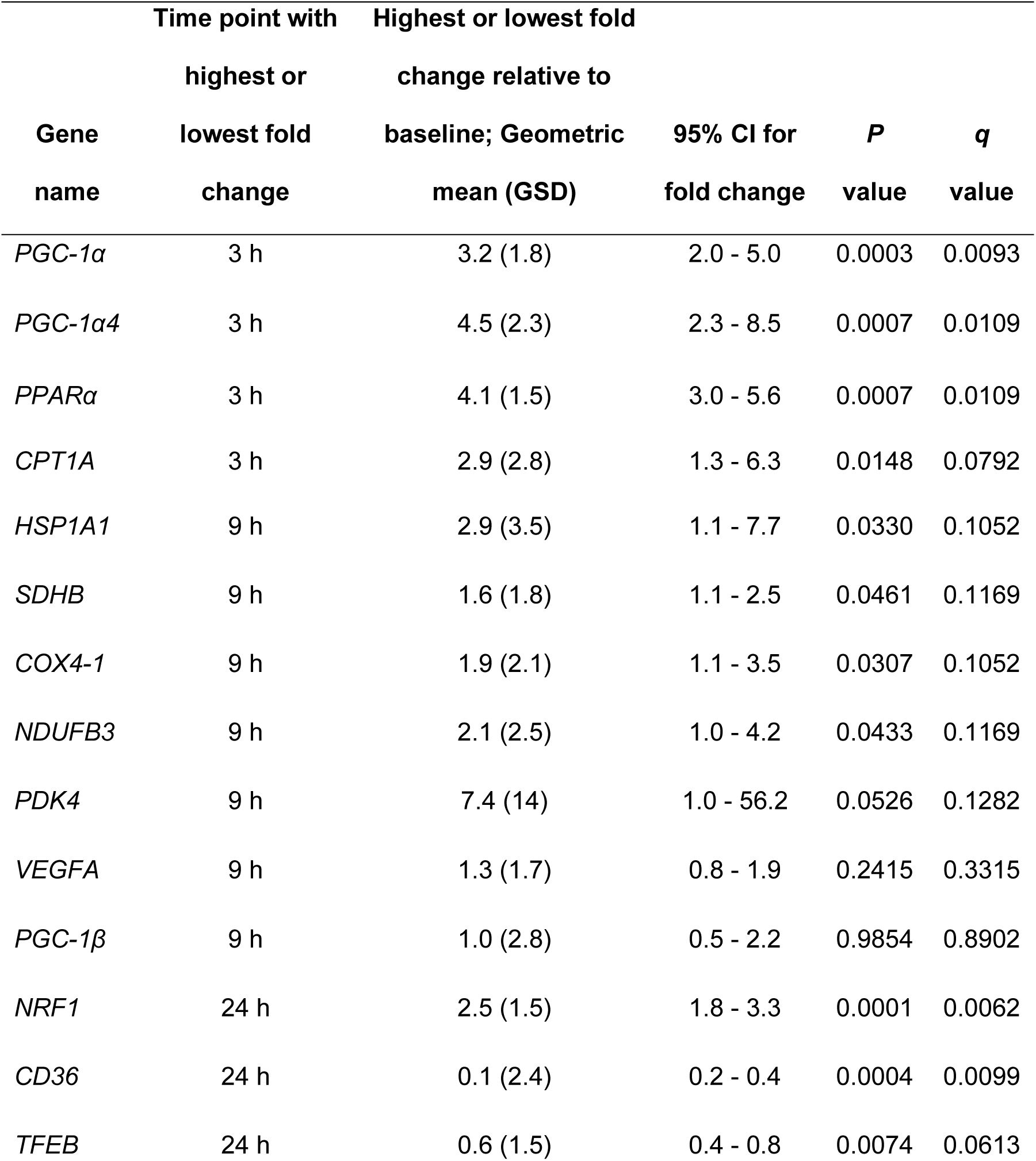
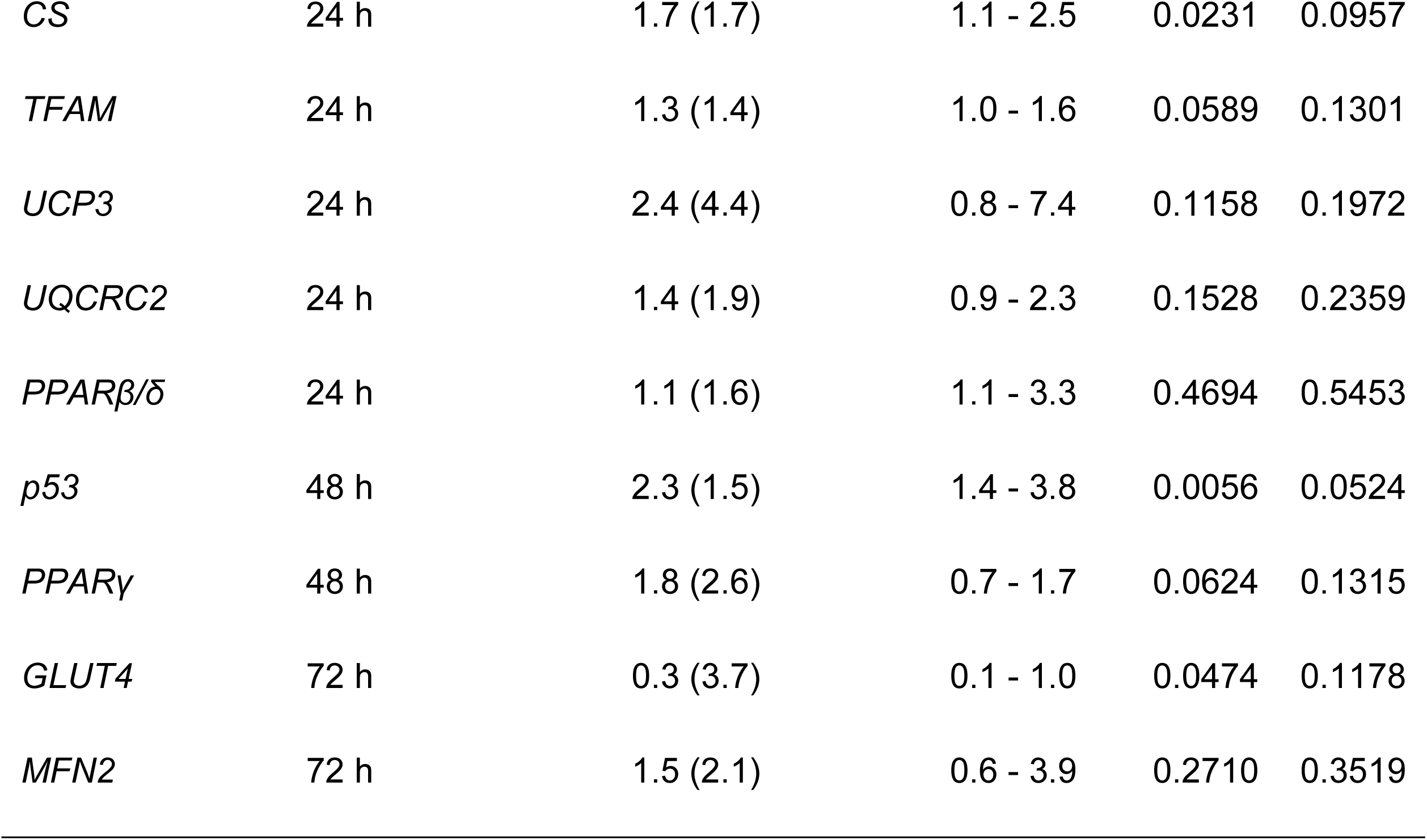
Summary of changes in mRNA content following a single session of HIIE, measured in nine participants. The time point with maximal fold changes, geometric mean for maximal fold change with geometric standard deviation (GSD), 95% Confidence Interval (CI), and *P* value and *q* value determined by a Benjamini-Hochberg false discovery rate (FDR) of < 10% are reported for each target gene.

**Table 2.** Function of genes that responded to a single session of HIIE.

| Gene | Main function in exercise adaptation<br>(NCBI Reference sequence database) |
| --- | --- |
| PGC-1α | This gene encodes a transcriptional coactivator that regulates the genes involved in energy metabolism as a transcription factor. It provides a direct link between external physiological stimuli and the regulation of mitochondrial biogenesis, and is a major factor that regulates muscle fibre type determination. |
| PPARα | This gene encodes the subtype PPAR-alpha, which is a nuclear transcription factor. PPARs affect the expression of target genes involved in cell proliferation, cell differentiation and in immune and inflammation responses. |
| HSPA1A<br>(HSP70) | This gene encodes a 70kDa heat shock protein which is a member of the heat shock protein 70 family. It is involved in stabilizing existing proteins against aggregation and mediating the folding of newly translated proteins. It is also involved in the ubiquitin-proteasome pathway. |
| SDHB | This gene codes the iron-sulfur subunit of the mitochondrial electron transport chain Complex II, which is specifically involved in the oxidation of succinate. |
| COX4-1 | This gene encodes a nuclear-encoded subunit of the terminal enzyme of the mitochondrial respiratory chain that couples the transfer of electrons from cytochrome c to molecular oxygen. |
| TFAM | This gene encodes a key mitochondrial transcription factor containing two high mobility group motifs. The encoded protein also functions in mitochondrial DNA replication and repair. |
| CS | This gene encodes a Krebs tricarboxylic acid cycle enzyme that catalyzes the synthesis of citrate from oxaloacetate and acetyl coenzyme A. |
| p53 | This gene encodes a tumor suppressor protein containing transcriptional activation, DNA binding, and oligomerization domains. The encoded protein responds to diverse cellular stresses to regulate expression of target genes, thereby inducing cell cycle arrest, apoptosis, senescence, DNA repair, or changes in metabolism. |
| GLUT4 | This gene encodes a protein that functions as an insulin-regulated facilitative glucose transporter. |

**Figure 1:**
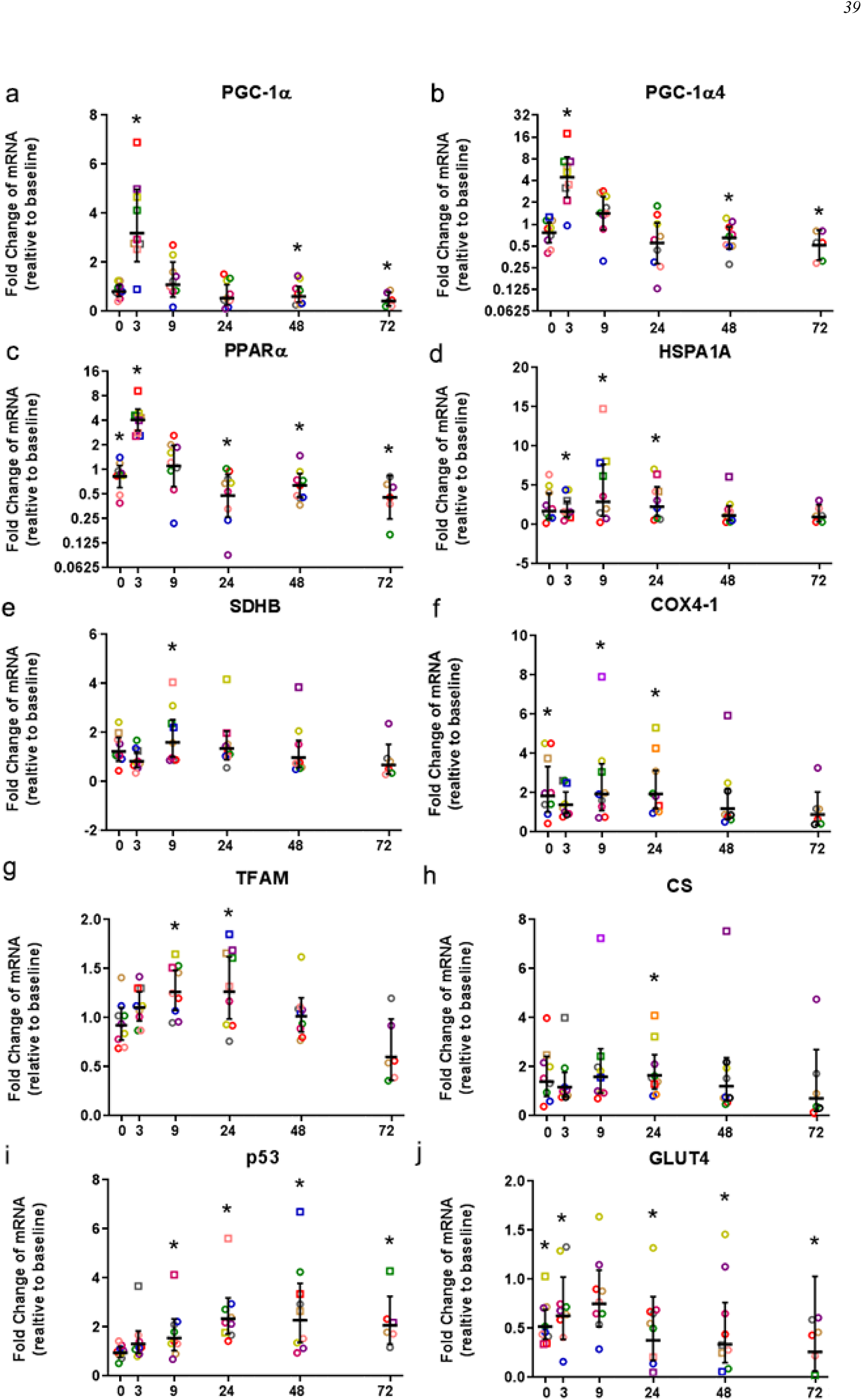
Relative fold changes compared to baseline for the mRNA content of *PGC-1α* (a), *PGC-1α4* (b), *PPARα* (c), *HSPA1A* (d), *SDHB* (e), *COX4-1* (f), *TFAM* (g), *CS* (h), *p53* (i), and *GLUT4* (j), following a single session of high-intensity interval exercise (HIIE). Muscle biopsies were obtained from nine participants at rest (baseline) before 4 weeks of High-Intensity Interval Training (HIIT), immediately after the final session of HIIE (0 h), and 3, 9, 24, 48, and 72 h after exercise, except only six participants at 72 h. Symbols (open circles and squares) of the same color indicate mRNA data from one participant; the geometric mean (horizontal bars) ± the 95% confidence interval (CI) are plotted for each graph. The squares indicate the data point with highest mRNA content for each participant. * Significantly different from baseline, determined by a Benjamini-Hochberg false discovery rate (FDR) of < 10%.

The geometric mean of mRNA content of *PGC-1α*, exercise-induced isoform *PGC-1α4*, and *PPARα*, increased significantly 3 h post exercise and was not significantly different from baseline at 9 h post exercise (Figures 1a to 1c). All nine participants had the highest expression level of *PGC-1α, PGC-1α4*, and *PPARα* mRNA at 3 h post exercise (except one participant who had the highest *PGC-1α4* mRNA content at 0 h). A similar result was observed for *CPT1A*; the highest mRNA level for the group geometric mean, and the highest value for seven out of nine individuals, occurred 3 h post exercise with values not significantly different from baseline at 9 h post exercise (data not shown).

The highest geometric mean for the mRNA content of *HSPA1A, SDHB*, and *COX4-1* occurred at 9 h post exercise (Figures 1d to 1f). The highest value for *NDUFB3* mRNA was also observed at 9 h. *PDK4, VEGFA*, and *PGC-1β* showed the greatest, but not significant, induction of mRNA content at 9 h post exercise (Table 1).

The geometric average mRNA content of *TFAM* and *CS* was highest at 24 h post exercise (Figures 1g and 1h), and most participants showed the highest mRNA content between 3 and 24 h post exercise (eight out of nine participants for *TFAM*, and seven out of nice participants for *CS*, respectively). *NRF1, CD36, TFEB, UCP3, UQCRC2*, and *PPARβ/δ* also showed the highest expression level at 24 h; however, only changes in *NRF1, CD36*, and *TFEB* reached significance (Table 1).

The geometric average *p53* mRNA level was significantly greater than baseline from 9 h to 72 h post exercise, with the highest expression 48 h post exercise (Figure 1i). *PPARγ s*howed the greatest, but not significant, induction of mRNA content at 48 h post exercise (Table 1).

Participants showed a small decrease of *GLUT4* mRNA content compared to baseline, and the lowest mRNA content was found at 72 h after exercise (Figure 1j). The decrease of the geometric average mRNA content was significant at all time points except 9 h post exercise. *MFN2* had the highest mRNA expression at 72 h post exercise; however, this change was not significant (Table 1).

### Modelling the gene expression response to exercise

We used a least-squares Gaussian nonlinear regression analysis to model the exercise-induced expression pattern of the target genes (Figure 2). Using the mRNA content from 0 to 48 h after exercise (the time span in which most of the transcriptional responses were observed), the best curve fit was generated based on the group mean mRNA content. The predicted peak mRNA expression time was identified based on the regression curve. The modelled peak mRNA expression time ranged from 4.6 h (*PGC-1α)* to 34.8 h (*p53*) post exercise (Figure 3). *GLUT4* was not used in this analysis as there was no peak of gene expression detected using curve fitting. We then established the time window within which each target gene expression level was within 90% of its maximal expression. The genes that responded earlier tended to have a shorter time window within 10% of maximal expression, whereas the genes that responded later had a larger time window.

**Figure 2:**
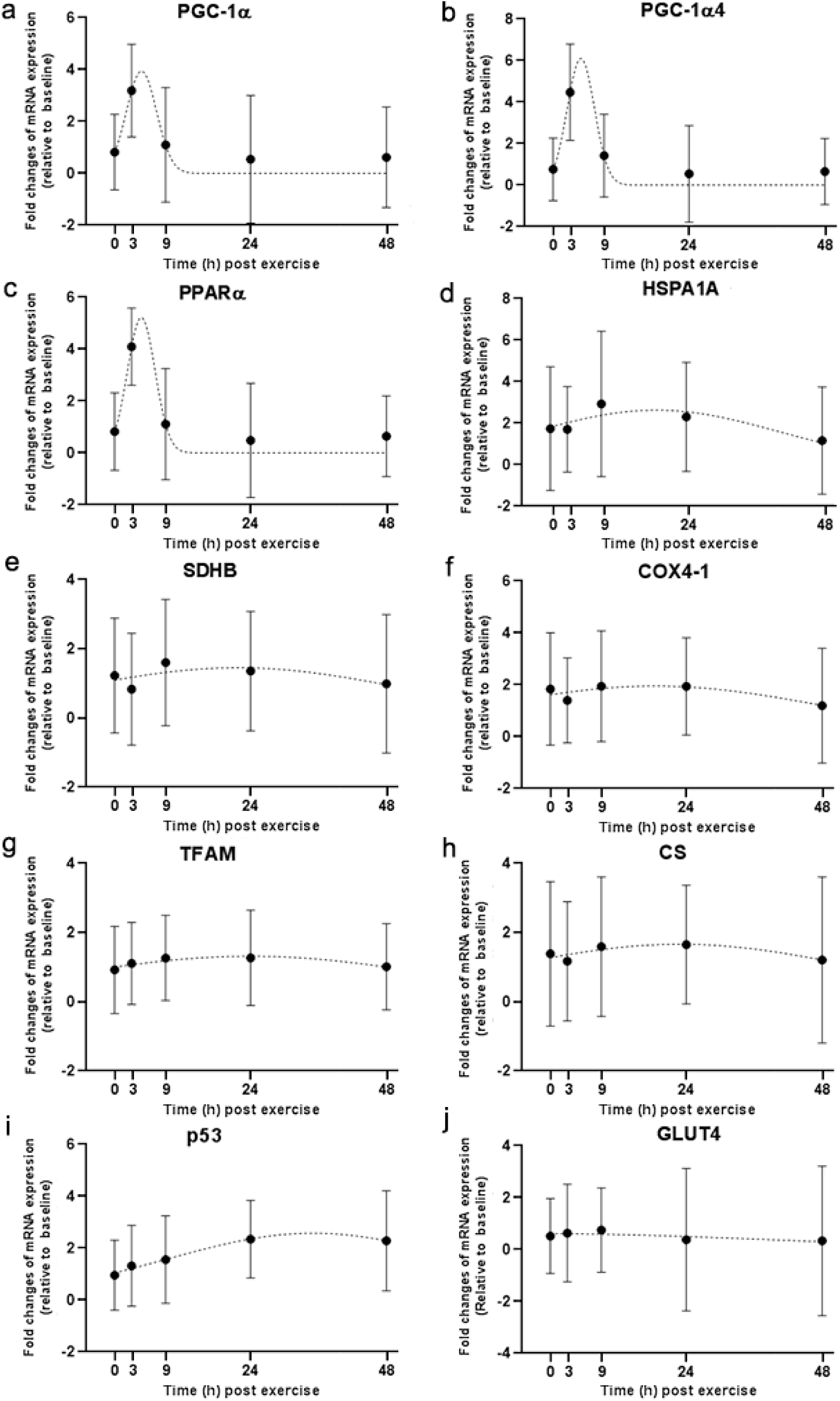
Curve fitting applied to mRNA changes following a single session of high-intensity interval exercise (HIIE). Least-squares Gaussian nonlinear regression analysis (dash lines) has been applied to gene expression data for *PGC-1α* (a), *PGC-1α4* (b), *PPARα* (c), *HSPA1A* (d), *SDHB* (e), *COX4-1* (f), *TFAM* (g), *CS* (h), *p53* (i), and *GLUT4* (j) at 5 time points (0, 3, 9, 24 and 48 h following a single session of HIIE). Geometric mean of gene expression is indicated by black dots, error bars are geometric standard deviations.

**Figure 3:**
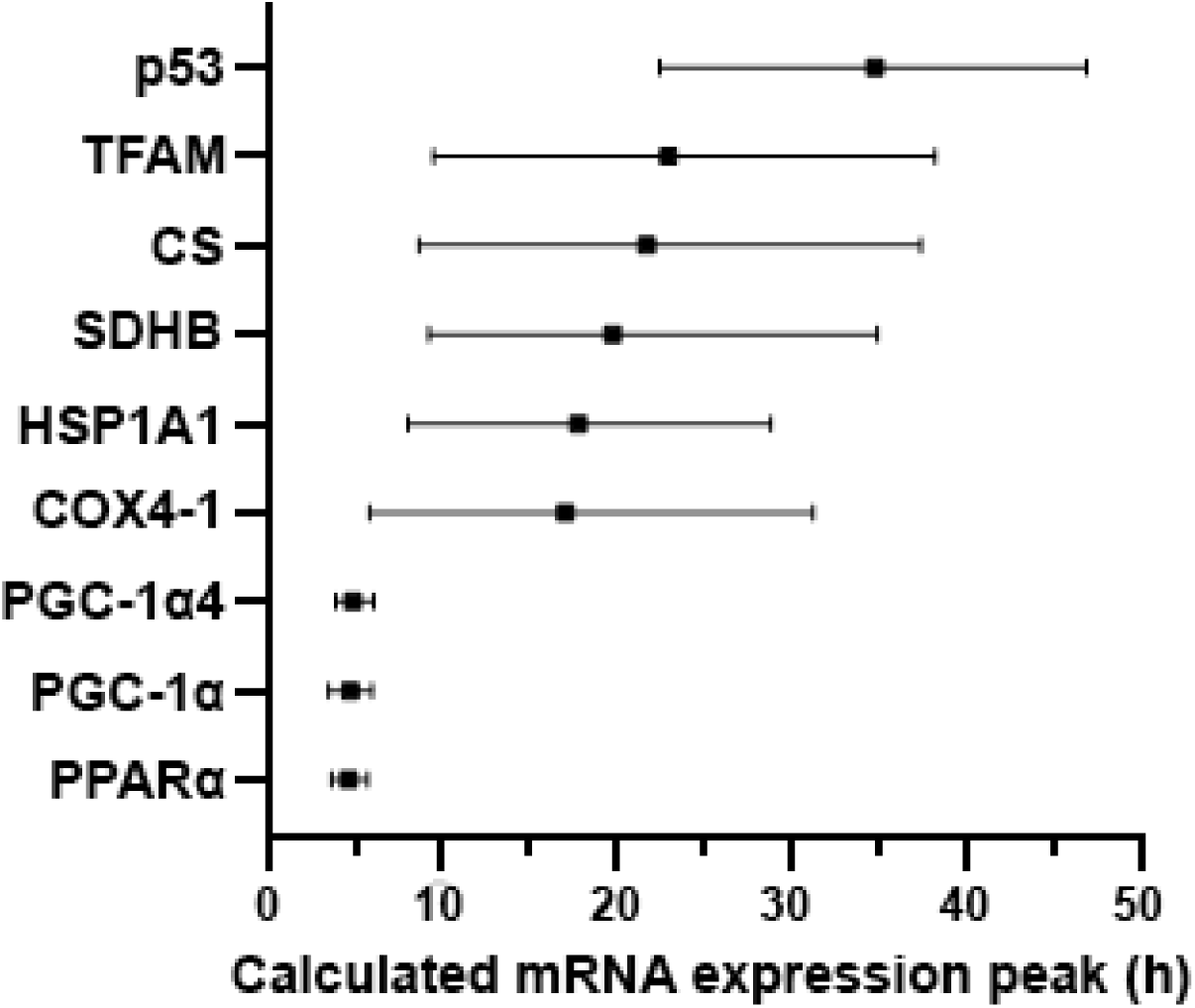
Modelled time of mRNA expression peak following a single session of high-intensity interval exercise in relation to biopsy timing. The peak expression time (black dots) and the time window for the top 10% of mRNA content (vertical lines) were calculated based on regression analysis and is shown for the nine genes for which a peak of gene expression was detected using curve fitting.

### Gene expression timing plotted against basal expression level

Using all 23 gene targets, we then plotted basal mRNA expression levels against the biopsy time that elicited peak mRNA expression after exercise. There was no clear relationship between the mRNA expression level at baseline and the biopsy time point that corresponded with the largest change in gene expression (Figure 4).

**Figure 4:**
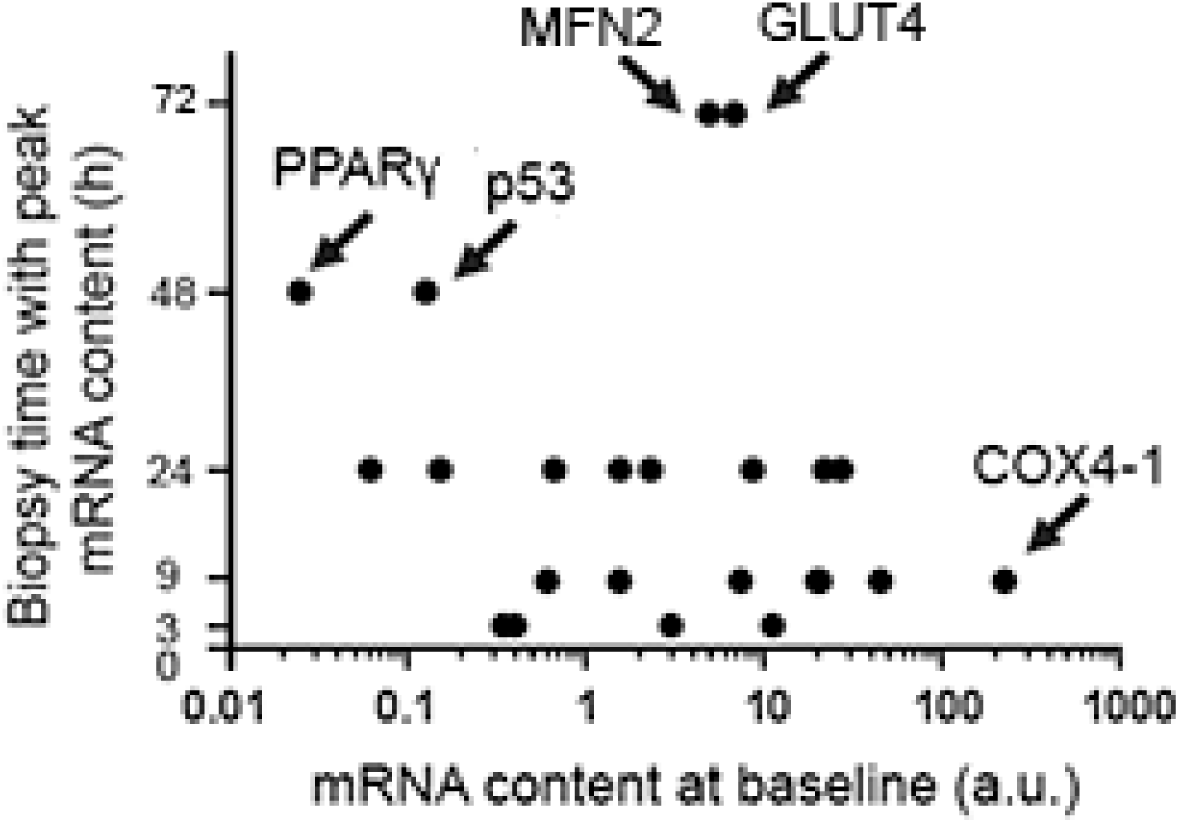
The biopsy time associated with peak mRNA expression plotted against the mRNA content at baseline for 23 gene isoforms.

## DISCUSSION

The present study examined changes in the mRNA content of 23 targets at six time points over a period of 72 h after a single session of high-intensity interval exercise in healthy participants. The changes in gene expression were highly dependent on the biopsy timing, and the greatest increase of mRNA content was observed between 3 h to 72 h post exercise. For many genes, there was considerable individual variability for the time point at which the maximal mRNA content was observed, especially for genes that responded later.

We observed distinct temporal patterns of gene expression after exercise, which is consistent with previous research. For example, Yang *et al.* ^23^ examined the mRNA content of several genes at seven time points from 0 h to 24 h after resistance exercise, and found that the timing of mRNA induction of their target genes was also variable. They reported that the mRNA levels of *muscle regulatory factor 4* (*MRF4*) and *PDK4* reached their maximal levels 4 h post exercise, whereas *myogenin* and *hexokinase II* (*HKII*) reached their maximal levels 8 h post exercise. Another study assessed exercise-induced gene expression at 3 h, 48 h, and 96 h after an endurance exercise session ^8^. Their results showed that the highest mRNA content of some targets, such as *hemeoxygenase 1* (*HMOX1*) and *integrin beta 2* (*ITGB2*), occurred 96 h post exercise, whereas other targets, such as *PGC-1α*, had the highest observed mRNA level at 3 h post exercise. These findings clearly highlight how biopsy timing has the potential to influence the interpretation of exercise-induced transcriptional responses. For example, if we had only taken a biopsy 3 h post exercise in the current study (which is common practice), we would have incorrectly concluded that genes such as *TFAM, HSPA1A, SDHB*, and *p53* were not affected by our exercise stimulus.

Genes that have a rapid and transient response after stimulation have been termed Immediate Early Genes (IEGs) ^30^. As the first responders at the transcriptional level, IEGs are required for progression to subsequent stages of transcription, such as the expression of subsequent “late response” genes ^31^. A subset of IEGs has been characterised as encoding for transcription factors ^30^. It is proposed that *PGC-1-related coactivator* (*PRC*), which is from the same family and shares structural and functional similarities with *PGC-1α*, belongs to the IEG class. This is consistent with our observation that the average mRNA content of *PGC-1α*, as well as exercise-induced isoform *PGC-1α4*, increased significantly 3 h after a single session of HIIE, and then returned to baseline at 9 h post exercise. Other studies have also reported that *PGC-1α* mRNA increases 2- to 15-fold, 2 to 5 h after a single session of exercise in humans (summarised in ^32^). In another review article, Islam *et al.* ^33^ compared 19 human studies with diverse exercise protocols and muscle sampling time points; all studies reported an increase in *PGC-1α* mRNA expression, except one study that performed a biopsy 24 h post exercise.

Compared with the rapid induction of IEGs, the induction of secondary response genes is delayed. It has been proposed this may be because it takes time for the translation of IEGs to transcription factors, which are required for the induction of secondary response genes ^34^. When we investigated downstream targets of PGC-1α protein (a transcriptional coactivator ^35^), we observed that the mRNA level of nuclear-encoded mitochondrial subunit *SDHB* was significantly higher at 9 h post exercise, and the mRNA content of *COX4-1* mRNA reached its highest level at 9 h post exercise. We also observed a later up-regulation of *TFAM* compared to *PGC-1α*, where most individuals had maximal mRNA expression between 9 and 24 h post exercise. Our results suggest that the downstream targets of master regulator PGC-1α are induced by a single session of exercise, but follow a delayed time-course. These findings support the proposal by Scribbans and colleagues that the absence of a systematic upregulation of PGC-1α targets could be because changes in some targets were not captured by their chosen biopsy time (3 h post exercise) ^28^. The review article by Islam *et al.* ^33^ also reported contrasting observations when the expression of downstream genes targeted by PGC-1α was examined, including *TFAM*. For 18 studies that observed an increase in *PGC-1α* mRNA level, 12 studies reported an increased expression of *TFAM* with at least one exercise protocol or time point (biopsy times ranging from 0 to 24 h, but mostly within 6 h post exercise), and six studies found no change (biopsy times ranging from 0 to 6 h post exercise). There is no clear explanation of the contrasting results; however, the authors suggested this lack of coordination could be due to the divergent temporal expression pattern of different genes, and recommended a more thorough investigation of gene expression time course in future human studies.

The half-life of mRNA is important for the kinetics of gene expression, as mRNA levels are determined by the rates of both RNA synthesis and degradation. It has been reported that many transcription factors and regulatory proteins have short half-lives ^36,37^. In a study that analysed mRNA half-lives in human B-cells, the authors reported that genes involved in transcription factor activity, transcription, and transcription factor binding are short-lived, with half-lives ranging between 3.6 to 4.3 h, whereas genes involved in glucose metabolism and the mitochondrial respiratory chain have longer half-lives of ∼10 h ^38^. This matches our observation that transcription factor and coactivators, such as *PGC-1α* and *PPARα*, are fast responding genes following exercise. In contrast, genes with functions in glucose metabolism (*GLUT4*) and the mitochondrial respiratory chain (*SDHB* and *COX4-1*) took longer to be induced after our exercise stimulus. Therefore, it is reasonable to suggest the diverse expression timing of different genes after exercise stimuli depends on the function/role of the target gene in the process of the adaptative response, rather than its basal expression level in skeletal muscle (i.e., time to peak expression did not seem to be related to mRNA expression level at baseline; Figure 4).

Due to limitations regarding the number of muscle biopsies that can be obtained in a single experiment, it was impossible to determine the precise peak expression time of each target gene in the present study. Thus, we decided to model the peak expression timing of target genes based on the observed mRNA changes in all participants. Previous time course studies in human subjects ^23,24,26,27^, as well as our present experiment, observed that the majority of the exercise responsive genes followed a similar pattern: an initial upregulation to an observed peak level followed by a return to baseline levels. Based on these observations, we chose least-squares Gaussian nonlinear regression modeling to analyse the expression pattern using the gene expression data from 0 to 48 h post exercise (as the observed maximal mRNA expression of all genes was within the first two days after exercise). Based on this model, we mapped out the time window eliciting the highest 10% of gene expression after exercise (Figure 3). One limitation of our modeling method, however, is that this parametric regression is strongly based on the assumption that the genes followed the modeled pattern. Furthermore, the large individual variation between participants, the relatively low number of participants, and the limited number of muscle biopsies taken, undoubtedly affected the accuracy of our regression analysis. An even more comprehensive study, with more participants, more biopsy time points, and a greater number of genes (assessed with microarray or RNAseq) is necessary to construct a more accurate picture of the kinetics of mRNA expression in response to exercise.

In order to standardise the exercise dose, we prescribed the same relative intensity and duration of HIIE for all participants. Nevertheless, we still observed large variability in both the timing and the magnitude of the transcriptional response between individuals. This is consistent with previous research showing that not all individuals respond the same way to a standardised exercise dose ^39-41^. This individual response could be due to genetic background, non-genetic biological and behavioral factors, day-to-day fluctuations, and technical or biological variability associated with the sampling of human skeletal muscle and the analysis of exercise-induced mRNA ^42,43^. We add that the timing of the observed peak for exercise-induced expression of different genes also differed between individuals. This further highlights that it is probably not possible to choose a single post-exercise biopsy time point that will capture changes in the expression of specific genes for all individuals.

It is also important to note that our results are likely to be specific for the participants recruited and the exercise employed. It has previously been reported that both training and fitness level can affect exercise-induced gene expression ^44,45^. Thus, the time courses we have reported cannot be assumed for other populations. Many other studies have also reported intensity-dependent changes in gene expression ^46-48^. It remains to be determined if the different time courses we have reported are also influenced by exercise intensity.

In summary, this study monitored the kinetics of mRNA expression with multiple time points over 72 h after a single exercise session. We observed distinct temporal patterns for the expression of different genes, with the time for the highest observed gene expression varying from 3 to 48 h post exercise. These findings highlight an important limitation when studying the molecular responses to exercise, where few (two to three) biopsy time points in a short time frame (less than 24 h) are commonly employed to examine transcriptional responses to exercise. These results further emphasise the importance of carefully planning biopsy time points to best capture changes in genes of interest and considering the biopsy time points chosen when interpreting any observed transcriptional results.

## MATERIALS AND METHODS

### Participants

As part of a larger project ^27,49^, sixteen recreationally-active men were fully briefed on the procedures, risks, and benefits associated with participating, before providing written informed consent to participate, and for their data to be used in the present study. However, due to the availability of muscle samples data from only nine participants were available for the present study, [mean (SD); age: 22 (4) y; height: 179.5 (7.9) cm; mass: 81.4 (14.3) kg; 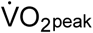: 3.9 (0.3) L•min^-1^]. All procedures were approved by the Victoria University Human Research Ethics Committee.

### Experimental Design

Following a familiarisation trial (performed on a separate day), participants completed a graded-incremental exercise test (GXT) to determine baseline levels of peak aerobic power 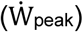 and the power at the first lactate threshold 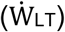. This information was used to individualise the intensity of a single session of high-intensity interval exercise (HIIE). As part of another study ^49^, the participants completed a 4-week high-intensity interval training (HIIT) intervention, training three days a week (12 sessions in total). The HIIE in the present study was the final HIIE session of the HIIT intervention. To avoid a response to muscle damage and unfamiliar stress ^50^, a resting muscle biopsy was taken before starting the 4-week training period (Week 0) and used as the baseline value, as done in previous research ^8,11^. Six more biopsies were sampled following the single HIIE session immediately (0 h), and 3, 9, 24, 48, and 72 h post-exercise (Figure 5).

**Figure 5:**
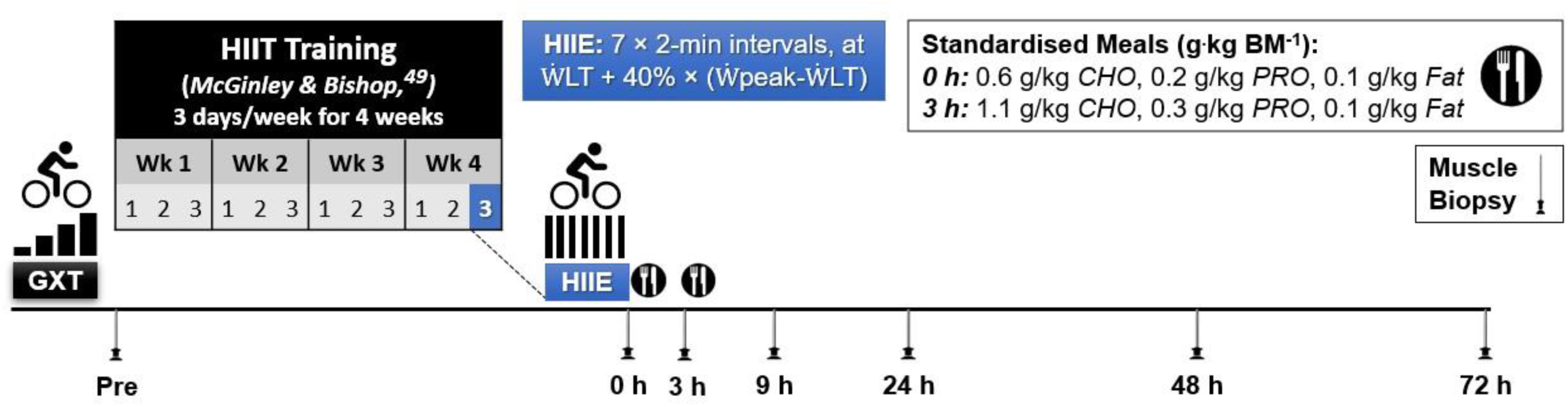
Experimental Design. Abbreviations: BM, body mass; CHO, carbohydrate; PRO, protein; GXT, graded exercise test; HIIE, high-intensity interval exercise; HIIT, high-intensity interval training; 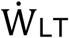, power at the first lactate threshold; and 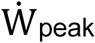, peak aerobic power.

### Graded Exercise Test

The GXT was conducted pre-training to determine the 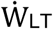 and 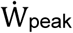. All trials were conducted in the morning (06.30–11.30). To determine the 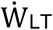, a 20-gauge intravenous cannula was inserted into an antecubital vein; venous blood was sampled at rest and at the end of every stage of the GXT, as previously described ^49^. The mean coefficient of variation (CV) for duplicate blood lactate measurements was 4.6%. The 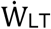 was identified as the power at which venous blood lactate increased 1 mM above baseline, and was calculated using Lactate-E software ^51^.

The GXT was performed on an electromagnetically-braked cycle ergometer (Excalibur Sport, Lode, Groningen, The Netherlands), using an intermittent protocol consisting of 4-min exercise stages separated by 30-s of rest ^52^. The initial load (90 to 150 W) was ascertained during the familiarisation GXT, and subsequently increased by 30 W every 4.5 min, with the aim of minimising the total number of stages to a maximum of ten. Participants were required to maintain a cadence of 70 rpm, and consistent verbal encouragement was provided during the latter stages. The test was terminated either volitionally by the participant, or by the assessors if the participant could no longer maintain the required cadence (± 10 rpm) despite strong verbal encouragement. 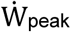 was calculated as previously reported ^53^.

### Peak Oxygen Uptake Test

After the GXT, the participants performed 5 min of active recovery at 20 W on the cycle ergometer, followed by a square-wave 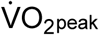 test. This comprised a steady-state cycle to volitional fatigue at a supramaximal power output, equating to the 105% of 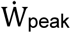 achieved during the GXT. A similar protocol has previously been reported to elicit 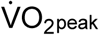 values no different to those determined during a ramp incremental test performed 5 min previously ^54^. The 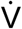 O_2_ peak was calculated as the mean of the two highest consecutive 15-s values. The exercise duration was 131 (37) s [mean (SD)].

### High-Intensity Interval Exercise

A single session of high-intensity interval exercise (HIIE) was performed between 06.30 and 08.00 following an overnight fast. The exercise consisted of seven 2-min intervals performed on an electromagnetically-braked cycle ergometer (Velotron, Racer-Mate, Seattle, WA), separated by 1 min of passive recovery (2:1 work:rest). A standardised 5-min steady-state warm-up at 75 W was completed beforehand. The exercise intensity was set to 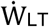 plus 40 % of the difference between 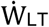 and 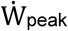; i.e., 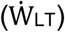 + (40 %)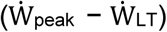. Power at the LT was 65 (6) % of 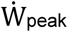, resulting in the HIIE bout being undertaken at 79 (4) % of 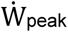 [mean (SD)]. Similar exercise protocols have previously been reported by our group to be effective at increasing 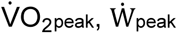, and 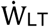 e.g. ^55,56-58^.

### Dietary Control

Participants performed the HIIE session following an overnight fast. Participants were requested to refrain from caffeine consumption on the day of the HIIE, to not ingest any dietary supplements, and to abstain from both alcohol consumption and exercise in the preceding 24-h period. Following the final HIIE session, participants were provided with two meals, totaling one-third of their daily energy requirement, based on their predicted basal metabolism ^59^, and allowing for a 1.4 activity correction factor ^60^.

Following the 0 h biopsy (i.e., immediately post-HIIE), participants received a standardised breakfast [1416 (230) kJ; mean (SD)], with a target macronutrient content (expressed in g per kg BM) of: 0.6 g/kg carbohydrate, 0.2 g/kg protein, and 0.1 g/kg fat. Following the 3 h biopsy, participants received a standardised meal [2456 (399) kJ], consisting of: 1.1 g/kg carbohydrate, 0.3 g/kg protein and 0.1 g/kg fat. The total relative macronutrient intake was therefore 64% carbohydrate, 16% protein and 20% fat. Excluding the standardised meals provided, participants were instructed to ingest only water *ad libitum* until after the 9 h biopsy. Light activities (e.g., walking) were permitted between the 0 and 9 h biopsies. All other biopsies (week 0 and 24, 48, and 72 h) were sampled in the morning following an overnight fast, with participants refraining from exercising or drinking alcohol until after the final muscle biopsy.

### Muscle Sampling

Muscle biopsies were taken from the belly of the vastus lateralis, approximately one third of the distance between knee and hip, using the needle biopsy technique modified with suction ^61^. Subsequent samples were taken approximately 1 cm proximal to the previous biopsy site. A 5 mm incision was made under local anesthesia (1% Lidocaine) and a muscle sample was taken using a Bergström needle ^62^. Samples were blotted on filter paper to remove blood, before being immediately snap-frozen in liquid nitrogen, and then stored at −80°C until subsequent analyses. Muscle samples were taken from the non-dominant leg at rest pre-HIIT (Week 0) and four weeks later after the final HIIE bout – immediately post-exercise (0 h), 3 h, 9 h, 24 h, 48 h, and 72 h post-exercise.

### RNA extraction

qPCR was performed on *n* = 9 at all time points, except for at 72 h (*n* = 6) using method estabilished by our group ^63^. An RNeasy Plus Universal Mini Kit was used to extract total RNA from approximately 20 mg of frozen muscle. Samples were homogenised using a QIAzol lysis reagent and a TissueLyser II (Qiagen, Valencia, USA). The instructions for the kit were modified slightly to increase RNA yield by replacing ethanol with 2-propanol and storing samples at -20°C for 2 h ^63^. Purification of RNA samples was performed according to kit instructions using a genomic DNA (gDNA) eliminator solution containing cetrimonium bromide. A BioPhotometer (Eppendorf AG, Hamburg, Germany), was used to determine both the concentration and purity of the RNA samples (based on the A_260_/A_280_ ratio). RNA integrity of all samples was measured using a Bio-Rad Experion microfluidic gel electrophoresis system (Experion RNA StdSens Analysis kit) and determined from the RNA quality indicator [RQI: 8.8 (0.5)]. RNA was stored at −80°C until reverse-transcription was performed.

### Reverse transcription

One µg of RNA, in a total reaction volume of 20 µL, was reverse-transcribed to cDNA using a Thermocycler (Bio-Rad, Hercules, USA) and iScript RT Supermix (Bio-Rad, Hercules, USA) as per the manufacturer’s instructions. Priming was performed at 25°C for 5 min and reverse transcription for 30 min at 42°C. All samples, including RT-negative controls, were run on the same plate. cDNA was stored at −20°C until subsequent analysis.

### qPCR

Relative mRNA expression was measured by qPCR (QuantStudio 7 Flex, Applied Biosystems, Foster City, USA) using SsoAdvanced Universal SYBR Green Supermix (Bio-Rad, Hercules, USA). Primers were designed using Primer-BLAST ^64^ to include all splice variants, and were purchased from Sigma-Aldrich (see Table 3 for primer details). All reactions were performed in duplicate on 384-well MicroAmp optical plates (Applied Biosystems, Foster City, USA) using an epMotion M5073 automated pipetting system (Eppendorf AG, Hamburg, Germany), which enabled all samples to be analysed for two genes on one plate. A total reaction volume of 5 µL contained 2 µL of diluted cDNA template, 2.5 µL of mastermix, and 0.3 µL of 5 µM or 15 µM primers. All assays were run for 10 min at 95°C, followed by 40 cycles of 15 s at 95°C and 60 s at 60°C. Of six potential reference genes assayed, the three most stably expressed were TBP, PPIA, and B2M. Primer specificity was confirmed by melting curve analysis. The expression of each target gene was normalised to the geometric mean of expression of the three reference genes ^65^, and using the 2^−ΔΔCq^ method ^66^. Amplification of no-template controls and no-RT controls on each plate resulted in either no detectable PCR product or a high C_q_.

**Table 3.**
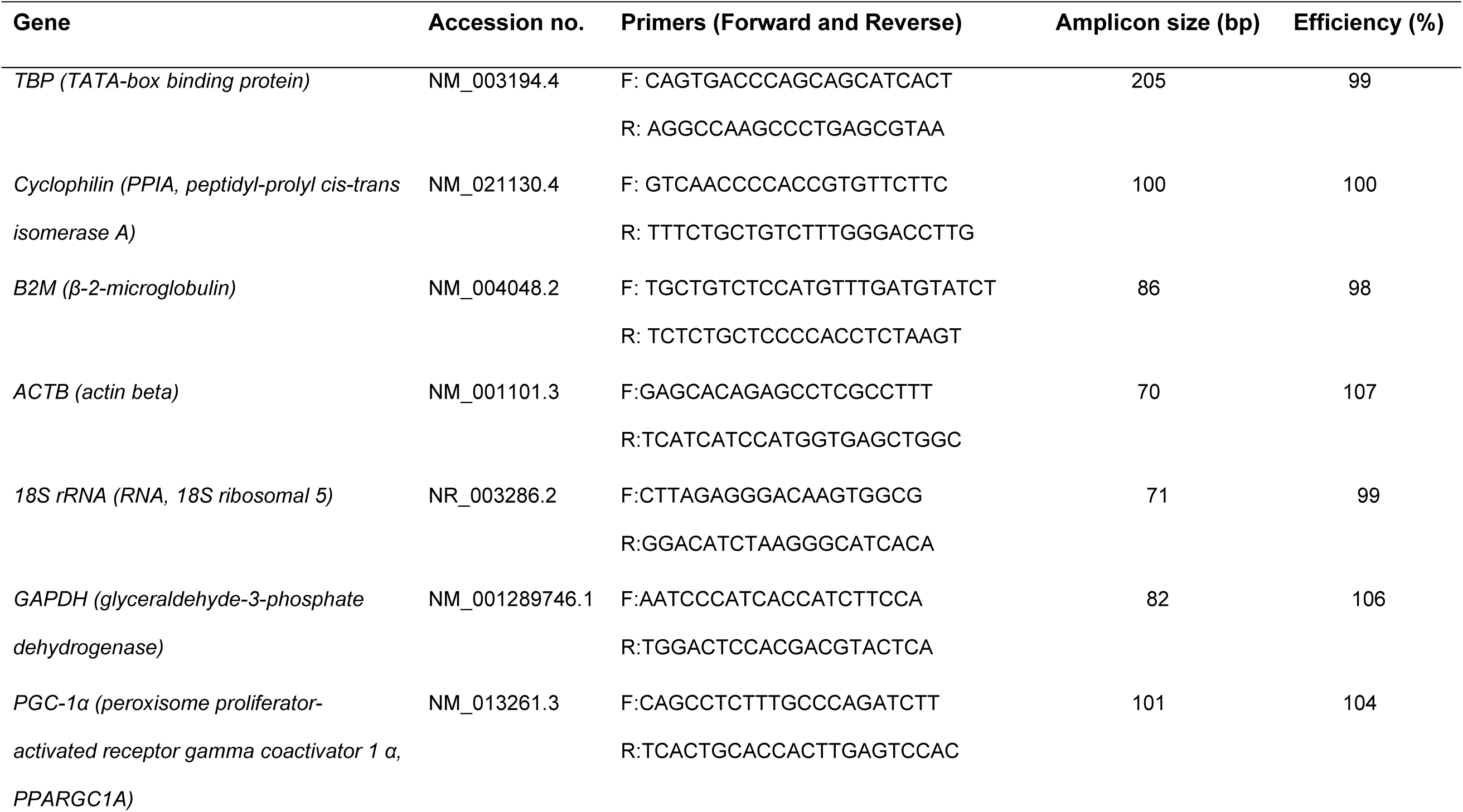
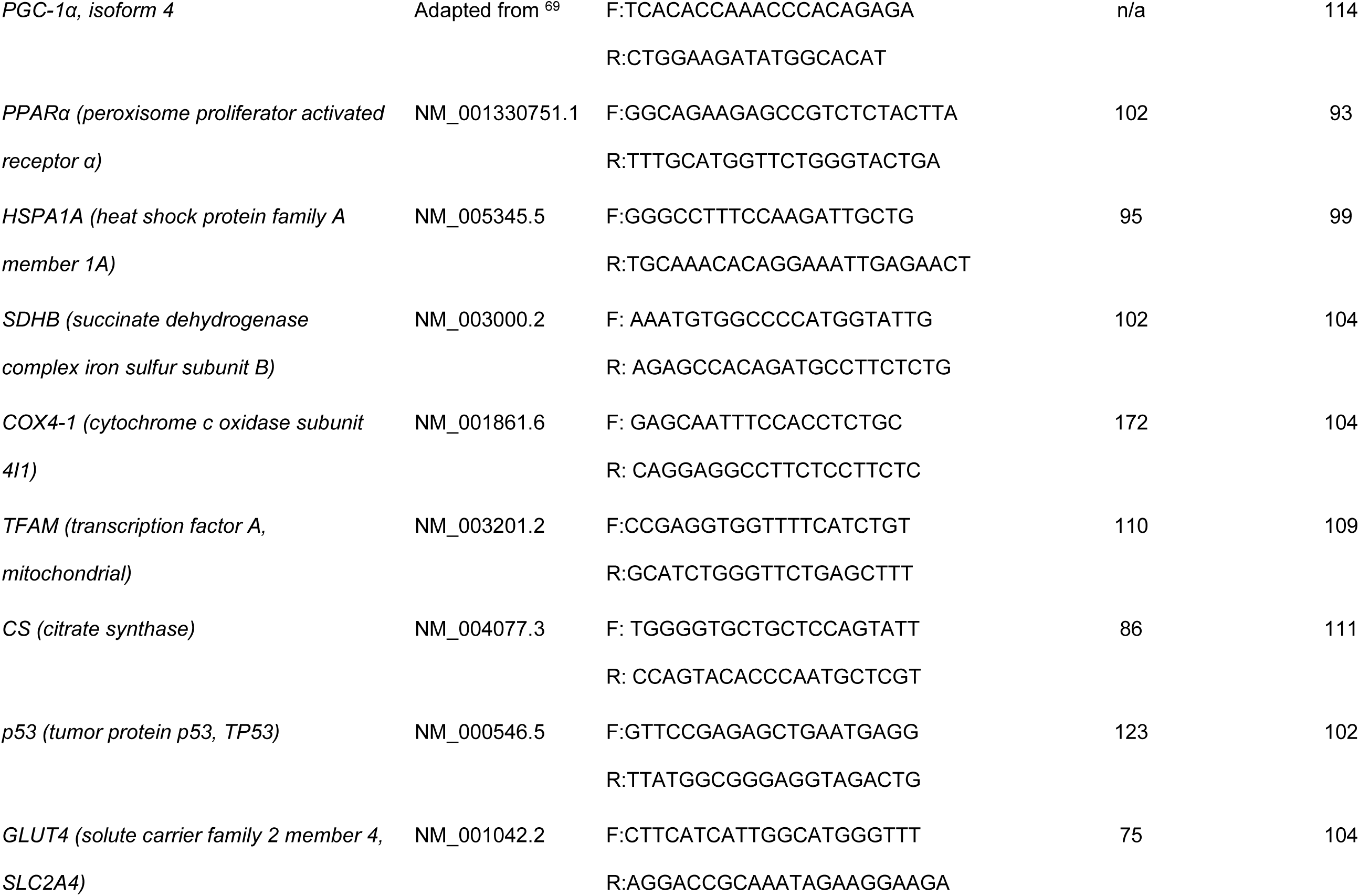
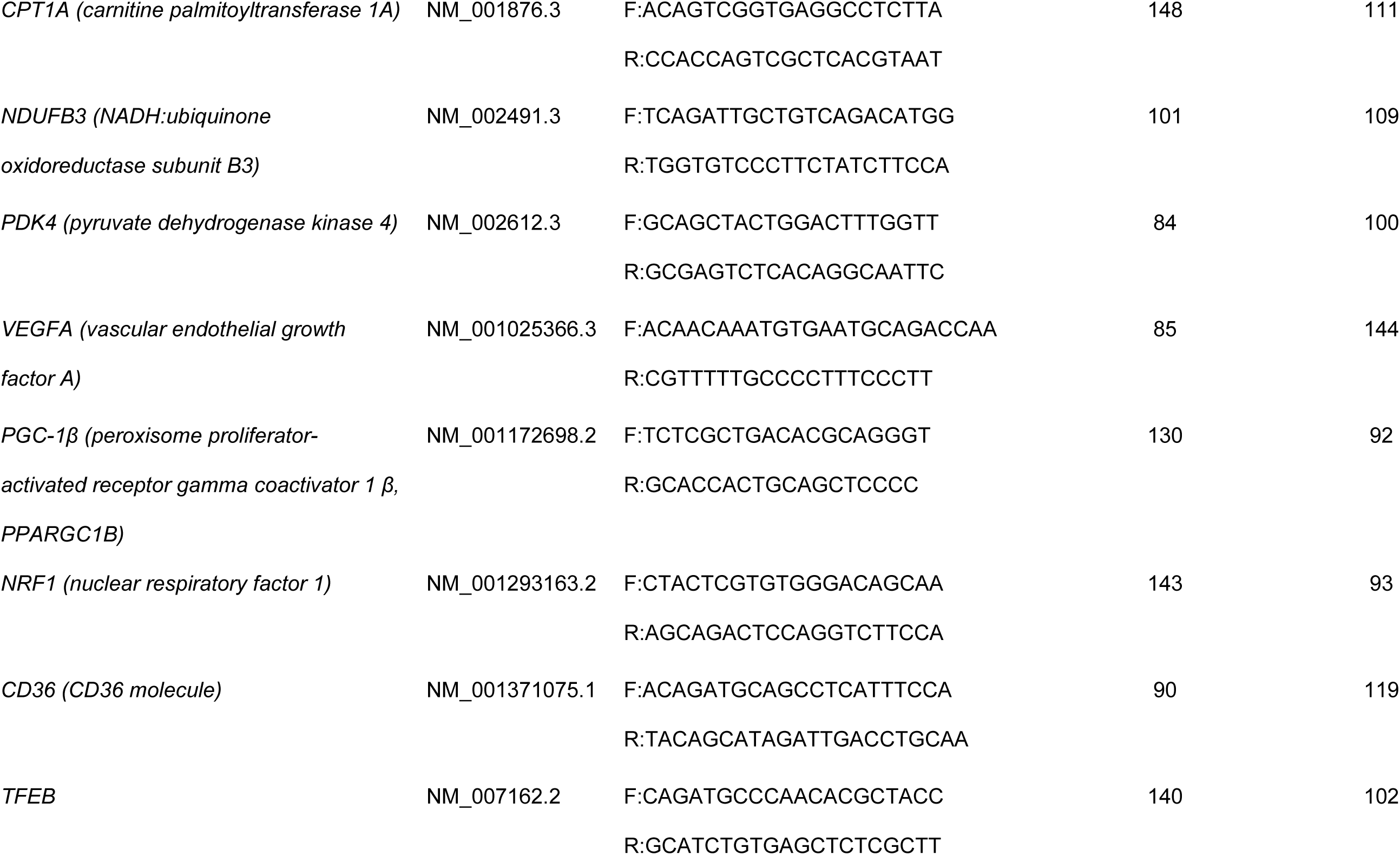
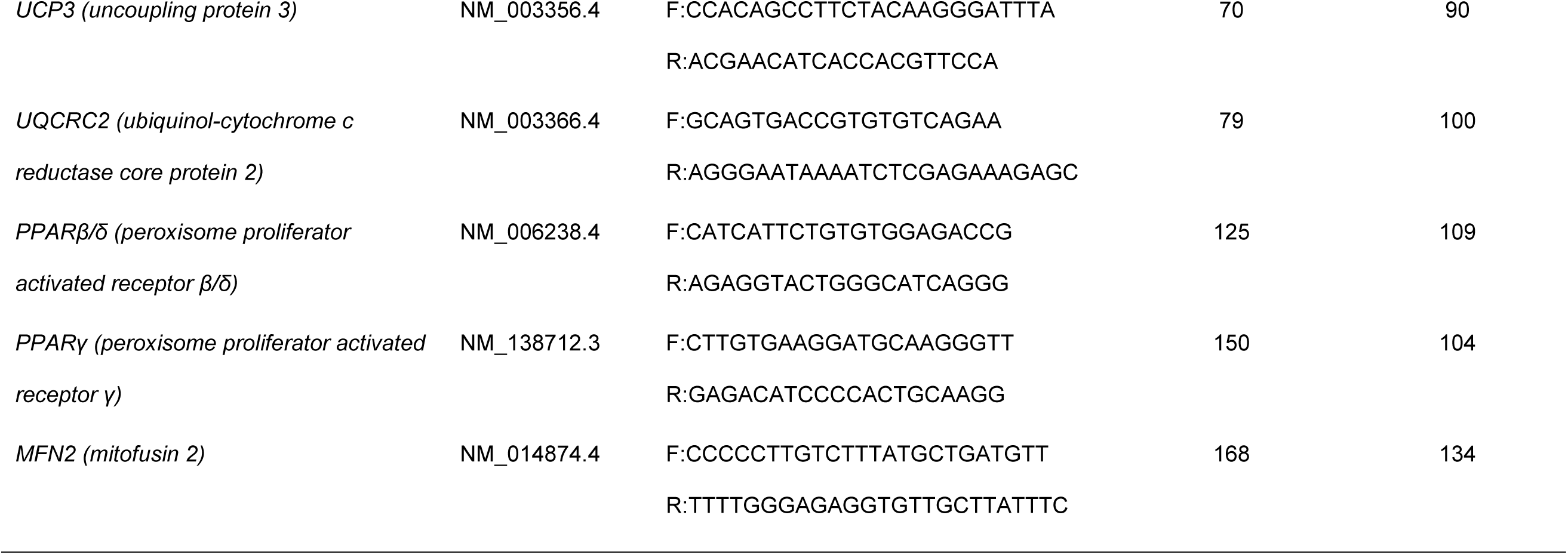
Primer sequences and amplicon details.

### Statistics

RefFinder was utilised for the statistical analysis of reference genes ^67^. For qPCR data present in Table 1, statistical analyses were performed on the log10 transformed 2^−ΔCq^ data, but the relative expression is reported (2^−ΔΔCq^). Geometric means and geometric standard deviations [geometric mean (GSD)] are reported. A paired t-test was used to compare the difference between time points post exercise and baseline values, and a Benjamini-Hochberg false discovery rate (FDR, Q) of < 10% was used to analyse all the *P* values (GraphPad Prism 8, GraphPad Software, Inc.). For data present in Figure 2, significance was determined by a Benjamini-Hochberg false discovery rate (FDR) of < 10% (GraphPad Prism 8, GraphPad Software, Inc.) ^68^. In Figure 3, the least-squares Gaussian nonlinear regression analysis (curve fitting) was used to model the peak of mRNA expression using GraphPad Prism 8 (GraphPad Software, Inc.).

## ACKNOWLEDGEMENTS

We are very grateful to the participants for their time and effort. We also thank Dr Mitch Anderson for performing some of the muscle biopsies.

## DATA SHARING

The data that support the findings of this study are available from the corresponding author upon reasonable request.

